# sRNA-target Prediction Organizing Tool (SPOT) integrates computational and experimental data to facilitate functional characterization of bacterial small RNAs

**DOI:** 10.1101/448696

**Authors:** Alisa M. King, Carin K. Vanderpool, Patrick H. Degnan

**Affiliations:** Department of Microbiology, University of Illinois, Urbana, IL 61801; Department of Microbiology and Plant Pathology, University of California, Riverside, Riverside, CA 92521

## Abstract

Small RNAs (sRNAs) post-transcriptionally regulate mRNA targets, typically under conditions of environmental stress. Although hundreds of sRNAs have been discovered in diverse bacterial genomes, most sRNAs remain uncharacterized, even in model organisms. Identification of mRNA targets directly regulated by sRNAs is rate-limiting for sRNA functional characterization. To address this, we developed a computational pipeline that we named SPOT for sRNA-target <u>P</u>rediction <u>O</u>rganizing <u>T</u>ool. SPOT incorporates existing computational tools to search for sRNA binding sites, allows filtering based on experimental data, and organizes the results into a standardized report. SPOT sensitivity (Correctly Predicted Targets/Total Known Targets) was equal to or exceeded any individual method when used on 12 characterized sRNAs. Using SPOT, we generated a set of target predictions for the sRNA RydC, which was previously shown to positively regulate *cfa* mRNA, encoding cyclopropane fatty acid synthase. SPOT identified *cfa* along with additional putative mRNA targets, which we then tested experimentally. Our results demonstrated that in addition to *cfa* mRNA, RydC also regulates *trpE* and *pheA* mRNAs, which encode aromatic amino acid biosynthesis enzymes. Our results suggest that SPOT can facilitate elucidation of sRNA target regulons to expand our understanding of the many regulatory roles played by bacterial sRNAs.

**IMPORTANCE:** Small RNAs (sRNAs) regulate gene expression in diverse bacteria by interacting with mRNAs to change their structure, stability or translation. Hundreds of sRNAs have been identified in bacteria, but characterization of their regulatory functions is limited by difficulty with sensitive and accurate identification of mRNA targets. Thus, new robust methods of bacterial sRNA target identification are in demand. Here, we describe our <u>S</u>mall RNA-target <u>P</u>rediction <u>O</u>rganizing <u>T</u>ool, which streamlines the process of sRNA target prediction by providing a single pipeline that combines available computational prediction tools with customizable results filtering based on experimental data. SPOT allows the user to rapidly produce a prioritized list of predicted sRNA-target mRNA interactions that serves as a basis for further experimental characterization. This tool will facilitate elucidation of sRNA regulons in bacteria, allowing new discoveries regarding the roles of sRNAs in bacterial stress responses and metabolic regulation.

## INTRODUCTION

Bacterial small RNAs (sRNAs) range in size from 30 to 300 nucleotides (nts). Regulation of mRNA targets by sRNAs via base-pairing dependent mechanisms alters translation or mRNA stability (1, 2). Most of the time, base-pairing interactions involve the 5’ or 3’ untranslated region (UTR) of the target mRNA but can also involve sites within the coding region of the target mRNA. Small RNA-dependent translational repression often occurs via interactions that directly interfere with ribosome binding to the mRNA. However, sRNAs have also been shown to activate mRNA targets through various mechanisms, including interference with mRNA decay (3, 4). In recent years it has become evident that sRNAs are ubiquitous and play an important role in mediating and regulating many basic cellular processes and stress responses. Hundreds of small RNAs have been identified in numerous bacterial species such as *Bacillus subtilis* (5), *Listeria monocytogenes*(6), and *Salmonella enterica* (7, 8). With the advancement of current technologies, the number of sRNAs identified in diverse organisms will surely increase. Consequently, there is a pressing need to develop new and better tools for sRNA characterization. In particular, there is a need for methods to address a major rate-limiting step in novel sRNA functional characterization, which is high-fidelity identification of mRNA targets.

A variety of computational and experimental methods have been used to predict and validate sRNA-mRNA target interactions. The computational tools currently available for sRNA target prediction, such as TargetRNA (9), sTarPicker (10), IntaRNA (11, 12), and CopraRNA (13), albeit powerful, have their limitations, the most problematic of which is the high rate of false positives. TargetRNA, sTarPicker, and IntaRNA all scan the entire genome and search for putative targets based on interaction hybridization energies. CopraRNA uses the same methodology as IntaRNA for predicting targets based on thermodynamic favorability of the interactions but goes a step further and also considers the conservation of those interactions across species, giving more weight to predictions that are conserved (13). When CopraRNA, IntaRNA, and TargetRNA were used in a side-by-side comparison, CopraRNA was found to have the highest positive predictive value (PPV) of 44% and reported the lowest rate of false-positives for known sRNAs across 18 enterobacterial species (14). Although CopraRNA possesses the highest PPV out of all tools, there were still substantial false positives reported. Moreover, CopraRNA is limited to identifying conserved sRNA-target RNA interactions and cannot identify species-specific interactions. As a result, caution should be used with these individual algorithms and they are frequently used in tandem with other target identification methods (14).

Experimental methods, including transcriptomic studies, have often been used to identify sRNA candidate targets. Transcriptomics methods uncover gene expression changes caused by absence or overproduction of an sRNA. While microarrays and RNA-sequencing have been successfully used to deduce sRNA targets, in many cases, separating direct effects from indirect effects is laborious and time-consuming. Moreover, the data obtained from transcriptomic studies can only reveal targets that are expressed under the specific growth conditions examined. As such, bona fide target genes that are poorly expressed or that are regulated by mechanisms that do not result in a substantial change in mRNA stability may be missed as sRNA targets. To address these issues, affinity purification methods have been developed to enhance identification of sRNA-mRNA interacting partners. For example, RIL-Seq (RNA interaction by ligation and sequencing) (15) identifies sRNA-mRNA partners that bind to the RNA chaperone Hfq (16) by co-immunoprecipitation, ligation, deep sequencing and analysis of RNA chimeras, which often represent true interacting partners. MAPS (MS2-affinity purification coupled with RNA-sequencing) (17) uses sRNA “bait” that is tagged with an MS2 aptamer and can be purified by interaction with the MS2 coat protein. RNA targets that are copurified with the sRNA bait are then identified by deep sequencing. Even with the variety of tools available for sRNA target identification, it is still not entirely clear which tools are the most effective for sRNA target identification.

In order to streamline the use of multiple existing sRNA prediction algorithms, we developed a software pipeline called SPOT (sRNA-target Prediction Organizing Tool) that uses several algorithms in parallel to search for sRNA-mRNA interactions. The software collates predictions and allows integration of experimental data using customizable results filters. First, we used two well-characterized *E. coli* sRNAs, SgrS (18) and RyhB (19, 20), to assess the effectiveness of SPOT as the targets of these sRNAs are well defined. Next, we extended the application of the SPOT pipeline to UTRs of mRNAs to identify potential sRNAs involved in regulation. We then applied the same parameters and analyses to a less characterized *E. coli* sRNA, RydC. Employing a combinatorial approach through SPOT predictions and experimental validation, we were able to identify two new RydC targets, *pheA* and *trpE*, which were downregulated and upregulated, respectively, by RydC.

## MATERIALS AND METHODS

### Software pipeline

A software pipeline was constructed in PERL to provide a single interface for running four sRNA-mRNA target prediction algorithms in parallel and collating their results (Fig. 1). Source codes for TargetRNA2 v2.01 (9), sTarPicker (10), IntaRNA v1.0.4 (12), and CopraRNA v 1.2.9 (13) were downloaded and installed on a multicore local server. The pipeline is comprised of 4 steps described briefly here.

1. Reference genome files are retrieved from RefSeq or local customized genome files can be used, provided they are in an appropriate RefSeq format (GBK file or PTT and FNA files).
2. Simultaneous searches are initiated for TargetRNA2, sTarPicker, and IntaRNA according to user defined search parameters (e.g., window size, seed size, significance cutoffs). Optionally, if RefSeq IDs and corresponding sRNA sequences from related genomes are provided, a CopraRNA search is initiated.
3. The pipeline tracks the progress of each job and once each search is completed the raw results files are read into memory.
4. User-defined results filtering parameters are applied (e.g., list with known binding coordinates, differential expression, operon data) and the raw results in memory are collated into a unified report.

**Figure. 1.**
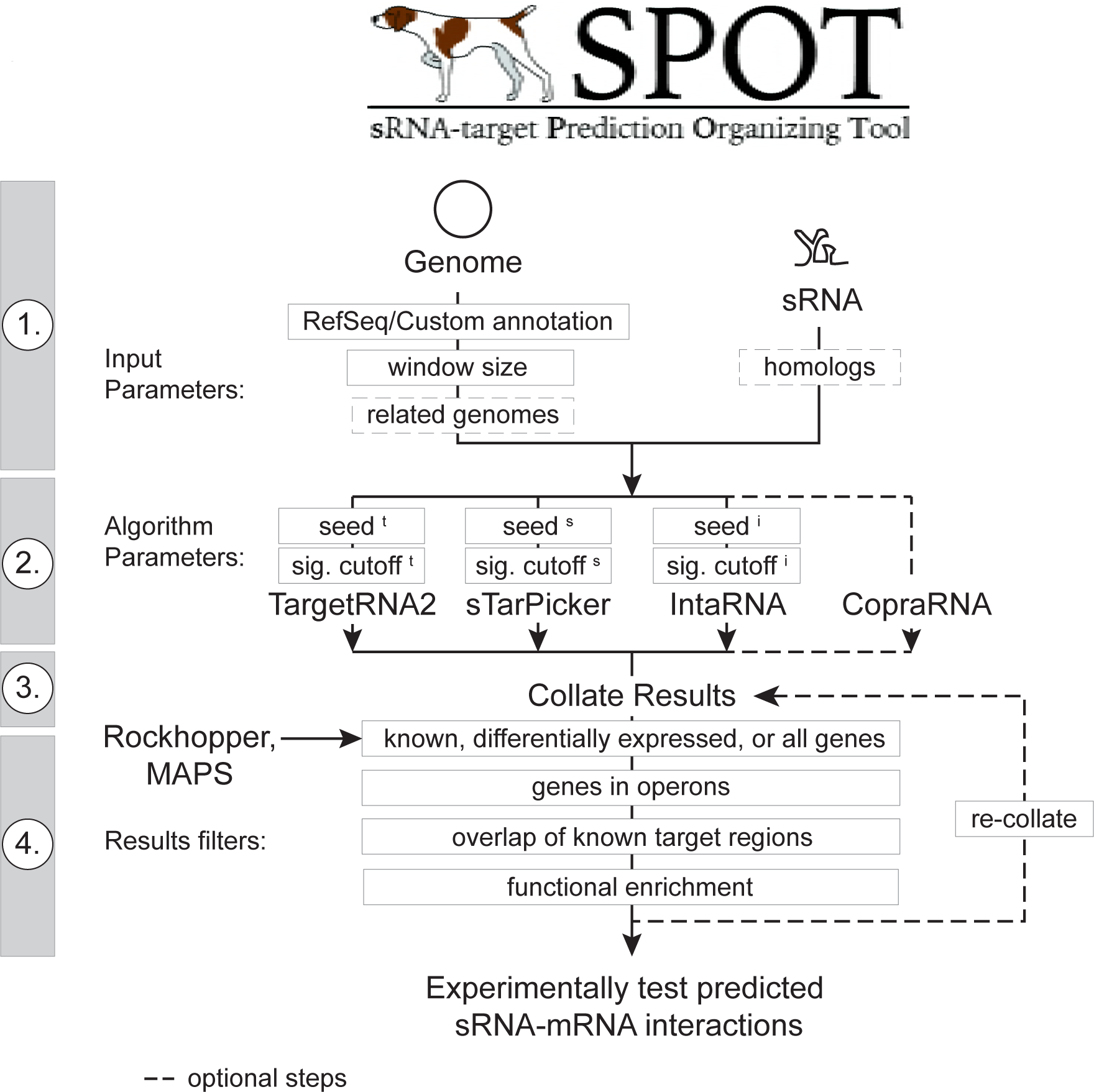
Schematic diagram of SPOT pipeline. Step 1: A basic implementation of SPOT requires a user-provided reference genome and a sRNA sequence file. The user can customize the search window size and can optionally provide information required for CopraRNA (dashed boxes). Step 2: The user can set seed sizes and significance cutoffs for each algorithm (superscript t = TargetRNA2, s = sTarPicker, i = IntaRNA). Step 3: SPOT runs the algorithms in parallel and generates a set of collated results. Step 4: Results filtering options as shown narrow the list of predicted interactions to an experimentally-tractable size for further validation or analysis.

The collated results report includes Excel-formatted data tables, functional enrichment predictions for consensus mRNA targets as well as binding plots. Both the collated results and individual search results can be downloaded once the job is complete. In addition, users can elect to have an email notification sent when the job is complete. The pipeline also includes an option to re-run the results collation steps using different results filters. This enables users to make minor adjustments to the results reporting without waiting for the individual searches to be re-run.

The SPOT program and installation instructions are available on GitHub (https://github.com/phdegnan/SPOT). In addition, an Amazon Web Service (AWS) cloud Amazon Machine Image (AMI) with all of the required software installed is available (search for SPOTv1). The SPOT User Manual is also included in Supplementary Material.

### Generation of test data sets

Known sRNA-mRNA interactions were collected from ecocyc.org (21), the literature, and experiments herein for 12 sRNAs with ≥4 confirmed targets: RyhB (b4451, RF00057), Spot42 (spf, b3864, RF00021), SgrS (b4577, RF00534), RybB (b4417, RF00110), FnrS (b4699, RF01796), GcvB (b4443, RF00022), OmrA (b4444, RF00079), CyaR (b4438, RF00112), MicA (b4442, RF00078), MicF (b4439, RF00033), DicF (b1574, RF00039), and RydC (b4597, RF00505) (Table S1). The confirmed sRNA-mRNA binding interactions were used as true positives, to investigate the reliability and sensitivity of the pipeline.

In order to test CopraRNA, homologs for the 12 *E. coli* MG1655 sRNAs were identified in related genomes using Infernal (22). For all sRNAs excluding DicF, the genomes of *E. fergusonii* ATCC 35469 (NC_011740), *Citrobacter koseri* ATCC BAA-895 (NC_009792), and *Salmonella enterica* sv. Typhimurium LT2 (NC_003197) were queried with the Infernal algorithm and each covariance model. For the sRNA DicF, a phylogenetically restricted sRNA, *E. coli* O157:H7 str. Sakai (NC_002695) and *E. coli* str. APEC O1 (NC_008563) were queried. In cases where genomes encoded ≥1 prediction (e.g., OmrA), the prediction with the lowest E value was used.

In addition, we compiled a list of 85 *E. coli* sRNAs to investigate the ability of the pipeline to be used to predict mRNA-sRNA interactions using a putative mRNA target as the search query (Table S2). This includes 65 RefSeq annotated sRNAs (NC_000913.3), an additional 19 sRNAs annotated in ecocyc.org (21), and the sRNA IepX (23). Note that 552 additional predicted *E. coli* sRNAs, *cis* regulatory elements and other putative RNAs corresponding to known RFAMs (n=172) or identified from expression studies (n=360) were not included (24, 25).

Finally, sRNA-mRNA interaction coordinates and the 5’ UTRs of 11 mRNAs with ≥2 known interacting sRNAs were collected from ecocyc.org (21): *csgD* (b1040,n=5), *flhD* (b1892,n=4), *ompA* (b0957,n=3), *ompC* (b2215,n=3), *ompF* (b0929,n=2), *ompX* (b0814,n=2), *phoP* (b1130,n=2), *rpoS* (b2741,n=4), *sdhC* (b0721,n=3), *sodB* (b1656,n=2), and *tsx* (b0411,n=2).

### Media and reagents

*E. coli* strains were cultured in lysogeny broth (LB) medium or on LB agar plates at 37°C, unless stated otherwise. For construction of reporter fusions by λ Red, recovery of recombinants was carried out on M63 minimal medium containing 5% sucrose, 0.001% L-arabinose (Ara), 0.2% glycerol, and 40 μg/ml 5-Bromo-4-Chloro-3-Indolyl β-D-Galactopyranoside (X-Gal). For β-galactosidase assays, bacterial cells were grown in Tryptone Broth (TB) medium supplemented with 100 μg/ml ampicillin (Amp) overnight at 37°C and then subcultured in TB broth containing 100 μg/ml ampicillin (Amp) with 0.002% L-arabinose. Where necessary, media were supplemented with antibiotics at the following concentrations: 100 μg/ml ampicillin (Amp), 25 μg/ml chloramphenicol (Cm), and 25 μg/ml kanamycin (Kan). Expression of RydC was induced with either 0.1 or 0.5 mM Isopropyl β-D-1-thiogalactopyranoside (IPTG) from the PLlacO-1 promoter.

### Strain construction

Strains and plasmids used in this study are listed in Table S3. All strains used in this study are derivatives of *E. coli* K12 strain MG1655. Oligonucleotide primers and 5’-biotinylated probes used in this study are listed in Table S4 and were all acquired from Integrated DNA Technologies (IDT). Chromosomal mutations were made by λ Red recombination (26, 27), and marked alleles were moved between strains by P1 *vir* transduction (28). PCR products were generated using the Expand^™^ High Fidelity PCR System (Sigma-Aldrich, St. Louis, MO) according to the manufacturer’s instructions. All mutations were verified by amplifying PCR fragments using GoTaq polymerase (Promega, Madison, WI) and sequencing.

The translational *lacZ* reporter fusions under the control of the PBAD promoter were constructed by PCR amplifying a fragment of interest using forward and reverse primers containing 5’ homologies to PBAD and *lacZ*(Table S3). PCR products were recombined into PM1205 using λ Red homologous recombination and counter-selection against *sacB* as described previously (29). The fusions used in this study were inserted into the *lac* locus of PM1205. Some *lacZ* reporter fusions used in this study were constructed using the one-step recombination method (30).

Plasmids harboring mutated *rydC* alleles under the control of the PLlacO-1 promoter were constructed using the Quickchange II XL Site-Directed Mutagenesis Kit (Agilent Technologies, Santa Clara, CA) with oligonucleotides AKP59 (PLlacO-1-*rydC3*), AKP68 (PLlacO-1-*rydC5*), and AKP69 (PLlacO-1-*rydC345*) that contained mismatched bases at the desired locations and transformed into XL10-Gold Ultracompetent cells (Table S3).

### RNA-seq analysis

*E. coli* K12 MG1655 strain AK250 (Δ*rydC, lacI ^q+^*) harboring vector (pBR322) or P_*lac*_-*rydC* plasmid was grown to OD_600_~0.5 in LB broth media at 37°C and then induced with 0.1 mM IPTG for 10 min. The hot phenol method (31) was used to extract total RNA after 2 and 10 minutes of induction. Samples were then treated with TURBO™DNase (Ambion) kit according to the manufacturer’s protocol and resolved by gel electrophoresis on 1.2 % agarose gel to confirm integrity of the 16S and 23S bands. Ribosomal RNA removal, library construction and sequencing were performed at the W. M. Keck Center for Comparative and Functional Genomics at the University of Illinois at Urbana-Champaign. Ribosomal RNA was removed from 1 μg of total RNA using Ribozero rRNA Removal Meta-Bacteria Kit (Illumina, Inc), and the mRNA-enriched fraction was converted to indexed RNA-seq libraries (single reads) with the TruSeq Stranded RNA Sample Preparation Kit (Illumina, Inc). The prepared libraries were then pooled in equi-molar concentrations and were quantified by qPCR with the Library Quantification kit Illumina compatible (Kapa Biosystems) and sequenced for 101 cycles plus seven cycles for the index read on a HiSeq2000 using TruSeq SBS version 3 reagents. The output fastq files were generated using Casava 1.8.2 (Illumina) and analyzed with Rockhopper (32). Genes were considered differentially expressed in RydC pulse-expression strains if they met a significance cutoff (q-value) of ≥0.005 and a fold-change value of >1.5 or <0.5. Some genes outside this range were studied because they met other criteria (e.g., prediction of a RydC-mRNA interaction by multiple algorithms).

### β-galactosidase assays

Bacterial strains were cultured overnight at 37°C (shaking) in TB medium containing 100 μg/ml Amp. Subsequent to overnight growth, cultures were diluted 1:100 into fresh TB media containing 100 μg/ml Amp and 0.002% Ara and cultured at 37 °C. After reaching an OD_600_ of 0.3, 0.1 or 0.5 mM IPTG was added to induce expression of the plasmids and grown for an additional hour until an OD_600_ of 0.5 - 0.6 was reached. All β-galactosidase assays were performed as described in previous protocols (33). In short, the samples were suspended in Z-buffer, with reactions conducted at 28°C with 4 mg/ml 2-nitrophenyl β-D-galactopyranoside (ONPG) as a substrate and 1 M Na_2_CO_3_ to end the reactions.

## RESULTS

### Integrated pipeline for sRNA target prediction algorithms

A number of algorithms and tools for identifying putative sRNA-mRNA interactions have been developed (9, 10, 12, 13). However, no single target prediction tool is 100% accurate, the tools implement distinct user-defined parameters, each tool uses a different format for reporting results, and tools are hosted on distinct web platforms. Our approach was to create a single pipeline incorporating existing computational tools to search for sRNA binding sites, producing a collated and standardized results report (Fig. 1). We incorporated the TargetRNA2 (9), sTarPicker (10), IntaRNA (12), and CopraRNA (13) tools into this pipeline because they are widely used and have open source code. Input for the pipeline minimally includes a fasta sequence for the sRNA and the RefSeq number for the target genome. Additional RefSeq genome IDs and homologous sRNA sequences can be provided if the user wishes to include CopraRNA results in the analysis. The pipeline interface also allows the user to define a set of parameters for the individual algorithms and results filters. In particular, the results can be filtered for genes with known binding sites or sets of genes that were identified as putative targets by experimental methods (e.g., RNA-seq, MAPS [MS2 affinity purification coupled with RNA sequencing] (17), RIL-seq [RNA interaction by ligation and sequencing] (15)). For instance, output from the RNA-seq analysis tool Rockhopper (32) can be used directly as a results filter. The program then follows four basic steps (1) download/validate input files, (2) simultaneously initiate computational tools, (3) track job progress and read individual raw results, (4) filter and collate results into a single report (Fig. 1). Finally, an option is provided that allows users to re-collate the results from an initial analysis using different results filter settings.

### Pipeline Optimization with SgrS and RyhB targets

SgrS and RyhB are two well-characterized model sRNAs in *E. coli* critical for glucose-phosphate (34) and iron limitation (35) stress responses, respectively. Numerous studies have confirmed 8 mRNA targets of SgrS (18, 36, 37, 38, 39) and 18 of RyhB (20, 40, 41, 42). We used these two sRNAs to test the utility and sensitivity of the pipeline. For RyhB, the entire 90-nt sequence was used as query for the bioinformatics search. For SgrS, only the 3’ 80-nts of the 227-nt sRNA was used as query, since this is the region involved in target RNA binding. Our initial optimization of the pipeline focused primarily on three parameters: ‘seed size’, ‘window size’ and ‘significance cutoffs.’ Each application utilizes distinct defaults for these parameters. For example, ‘seed size,’ defined as the number of contiguous base pairing interactions required to define an sRNA-mRNA match is set to a default value of 7 in TargetRNA2 and IntaRNA and 5 in sTarPicker. We varied the seed sizes for each algorithm and determined how different seed sizes impact the sensitivity of detection of true targets for SgrS and RyhB. Sensitivity is defined as Correctly Predicted Targets/Total Known Targets (i.e., true positive rate). For TargetRNA2, a seed size of 7 gave the highest sensitivity for correct target predictions, with 38% and 56% correct predictions, for SgrS and RyhB, respectively (Fig. 2A). For sTarPicker, the seed size giving the optimal sensitivity was 6, with 63% and 72% of known binding interactions identified for SgrS and RyhB, respectively. IntaRNA yielded the highest sensitivity of all three algorithms, again at a seed size of 6. IntaRNA correctly identified 100% of known SgrS interactions and 94% of known RyhB interactions (Fig. 2A). Based on these results, we used seed size settings of 7 for TargetRNA2 and 6 for IntaRNA and sTarPicker for all other analyses.

**Figure. 2.**
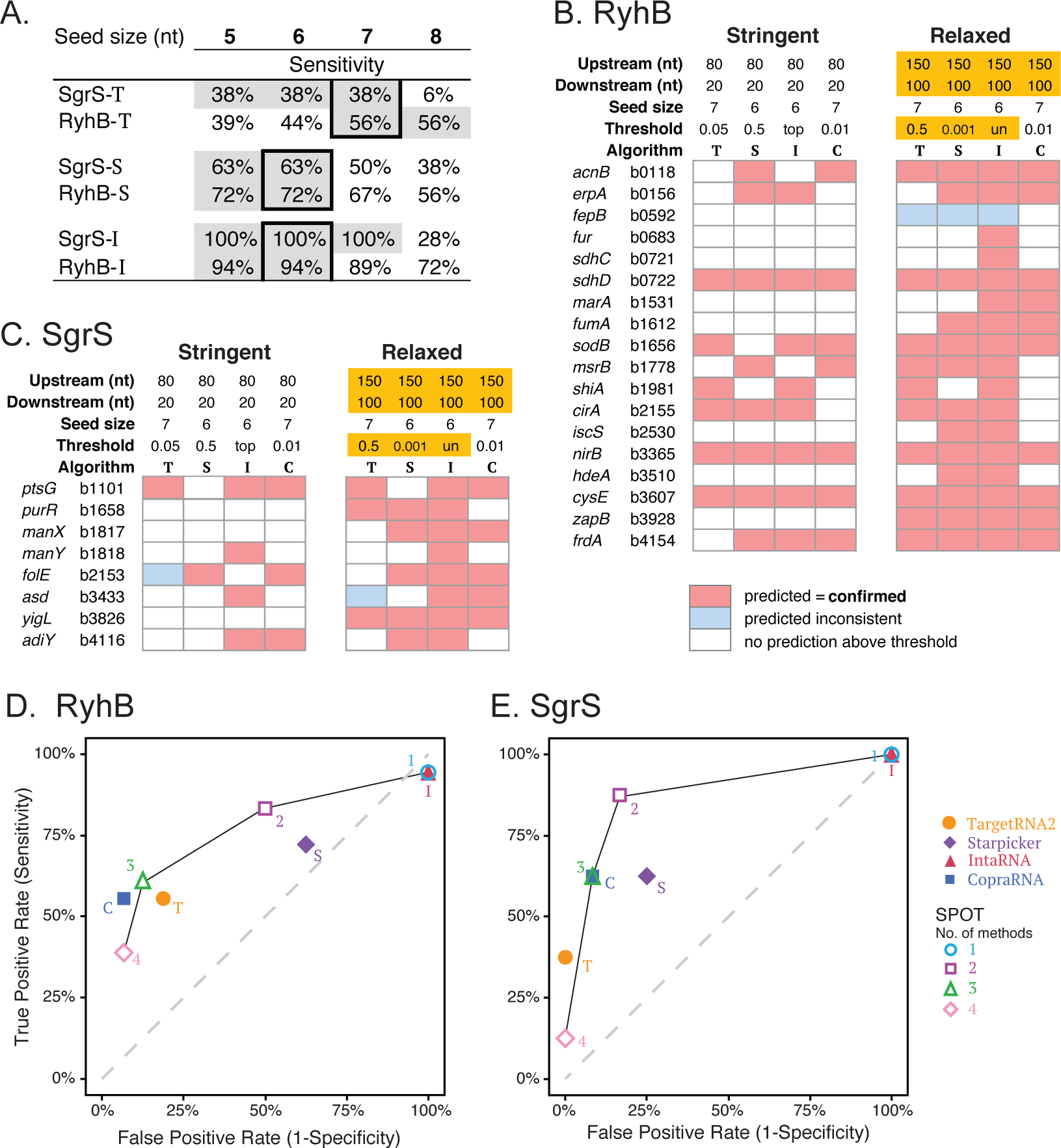
Validation of SPOT using known SgrS and RyhB sRNA-mRNA interactions. (A) “Seed size” indicates the number of consecutive basepairing nucleotides in an sRNA-mRNA interaction prediction. This is an adjustable parameter for each algorithm. Seed sizes were varied from 5 to 8 nt and the sensitivity (true positive rate) was determined for known SgrS and RyhB interactions while all other parameters were kept constant. Optimal seed sizes (bold boxes) were chosen for each algorithm. Highest percentage values for sensitivity are indicated with gray shading. Algorithms were abbreviated -T=TargetRNA2, -S=Starpicker, -I=IntaRNA. (B & C) Analyses were re-run using optimal seed sizes identified in A, but using a ‘Stringent’ parameter set with a narrow window size and high individual significance thresholds or a ‘Relaxed’ parameter set with a wider window size and low individual significance thresholds. Correctly predicted interactions for RyhB and SgrS are shown as pink cells, predictions that were inconsistent with confirmed interaction sites are shown in blue, and empty cells did not have any predictions above the indicated thresholds. Algorithms are abbreviated T=TargetRNA2, S=Starpicker, I=IntaRNA, and C=CopraRNA. (D & E) RyhB and SgrS have experimentally validated (true positive) and invalidated (true negative) mRNA targets, which were used to generate receiver operating characteristic (ROC) curves. These plots enable assessment of the accuracy of SPOT and the individual algorithms. Using the ‘Relaxed’ search parameters, 2-algorithm agreement in SPOT had greater True Positive Rates and more acceptable False Positive Rates compared with individual algorithms with the same settings.

Next, we evaluated how altering the window size and significance cutoffs impacted the accuracy of predictions (Fig. 2B, C). The window size refers to the size of the region upstream and downstream of every start codon in the genome that is searched for potential base pairing with the query sRNA. Default window sizes for each tool vary dramatically. The default TargetRNA2 window size is 80 nt upstream and 20 nt downstream (80/20) of each start codon (9). The default for sTarPicker is 150/100, IntaRNA suggests 75/75 and CopraRNA uses 200/100. Likewise, the tools have different metrics to determine the significance of a match either providing a *P* value (TargetRNA2, IntaRNA, CopraRNA) or a probability measure (STarPicker). TargetRNA2 generates *P* values for predicted interactions based on the sRNA-mRNA hybridization energy scores of a randomized mRNA pool (32). IntaRNA utilizes *P* values based on transformation of the energy scores calculated for all putative target binding sites with energy score ≤0 (14). CopraRNA combines individual IntaRNA *P* value predictions among clusters of genes to generate a weighted *P* value and false discovery rate (FDR)-corrected *Q* value (14). In contrast, sTarPicker uses a machine learning approach to generate probabilities as a proportion of base classifiers (n=1000) that support each proposed interaction (10). The sTarPicker authors report that probabilities ≥ 0.5 correspond to likely sRNA-mRNA interactions. SPOT provides the user with the ability to alter the search window and significance thresholds used by all the algorithms included in the pipeline (Fig. 1). We chose two sets of parameters that we define as “Stringent” and “Relaxed,” and tested the performance of each set of parameters in correctly identifying known RyhB and SgrS target binding sites (Fig. 2B, C). Stringent parameters incorporated a window size of 80-nt upstream and 20-nt downstream (80/20) of start codons as the search region, and significance thresholds of 0.05 for TargetRNA2, 0.5 for sTarPicker, “top” (e.g., the top 100 predictions) for IntaRNA, and 0.01 for CopraRNA (Fig. 2B, C). Relaxed parameters used a comparatively larger window size of 150/100 and thresholds of 0.5, 0.001, “un,” (e.g., all predictions) and 0.01 for TargetRNA2, sTarPicker, IntaRNA, and CopraRNA, respectively.

Using stringent search parameters, 10/18 known RyhB target binding sites and 2/8 known SgrS target binding sites were correctly predicted by ≥2 algorithms (Figure 2B, C, indicated by 2 or more pink cells and absence of blue cells). Using relaxed parameters, the correctly predicted interactions rose to 17/18 and 6/8 for RyhB and SgrS, respectively. Thus, for both RyhB and SgrS, relaxed parameters substantially increased the number of correctly identified binding sites (Fig. 2B, C). Notably, use of relaxed parameters was necessary to capture true binding sites like the SgrS binding site on *yigL* mRNA, which is located further from the start codon than is typical. The relaxed parameters improve the sensitivity of individual methods but may result in the downside of identifying more false positives. IntaRNA has high sensitivity for true positives (correct identification of known sRNA binding sites) under the relaxed settings, but also gives a high rate of likely false positives, illustrated by the fact that IntaRNA predicts >3400 binding interactions that are not predicted by any other algorithm. Mitigating this downside of using relaxed parameters, we saw that in the majority of instances the correct RyhB and SgrS binding sites were predicted by ≥2 methods and incorrect predictions by ≥2 methods occurred rarely (RyhB= 1/18, SgrS= 0/8) (Fig. 2B, C).

For SgrS and RyhB, at least a dozen mRNAs have been experimentally defined as ‘non-targets’ for each sRNA (18). In other words, predicted sRNA-mRNA interactions were tested and shown <u>not</u> to mediate regulation of the mRNA in question. These examples served as controls that allowed us to calculate False Positive Rates. Together with the Sensitivity measures for each algorithm and the pipeline, we generated receiver operating characteristic (ROC) curves to assess the accuracy of the methods alone and in combination (Fig. 2D, E). Ideally tools should yield high true positive rates and low false positive rates, resulting in values falling in the upper left quadrant of the ROC curve. Our results indicate that when 2 methods converge on the same prediction, the pipeline achieves ≥75% sensitivity and ≤ 50% false positive rate for both sRNAs. This is a marked improvement in most instances over the single algorithms used here (Fig. 2D, E). In particular, using a 2-method threshold mitigates the very high false positive rate from IntaRNA. We note that making the IntaRNA *P* value cutoff more stringent (e.g., 0.05) decreases the false positive rate dramatically, but at a cost to sensitivity (Fig. S1). Similarly, requiring 3 or 4 algorithms to identify the same predicted interaction decreases the false positive rate of predictions for RyhB and SgrS, however, the sensitivity decreases by more than 25% (Fig. 2D, E). Collectively, these analyses suggest that use of relaxed search parameters and a combined evidence approach requiring a minimum of 2 algorithms to predict the same binding interaction is an effective means of improving sRNA target prediction sensitivity.

The SPOT pipeline accepts several results filters to facilitate analysis of the predictions. First, users can provide the program a list of binding site locations for known mRNA targets (e.g., true positives). Second, users can include genes on the list that lack known binding sites in order to limit the results reporting to select genes of interest, for example those that emerged from experimental analyses (e.g., RNA-seq). Integration of experimental data with computational predictions is another valuable way of reducing potential false positive predictions.

Based on our results and observations during the optimization of SgrS and RyhB target identification, we designed SPOT to prioritize the target binding site predictions (Fig. S2). First, known binding sites correctly predicted by ≥2 algorithms (1) or 1 algorithm (2) are reported. Any gene targets with predictions that are discordant with known binding sites (3) are reported next. Then any additional targets with the same predicted target site found by ≥2 algorithms are ranked next (4). This is followed by targets that were only predicted by a single algorithm, in the following order: CopraRNA (5), TargetRNA, sTarPicker (6), and IntaRNA (7). Using the results filters, a user can narrow or widen their searches, for example, by limiting the predictions made by single algorithms or by applying secondary filters on binding site regions.

### Application of SPOT to additional sRNAs

To evaluate the robustness of the defined pipeline parameters and our ranking methods, we ran similar analyses on 9 additional sRNAs with ≥4 known targets. Overall, we found that the SPOT pipeline sensitivity (e.g., the percent of correctly identified interactions) was equal to or exceeded any individual method (average = 84% ± 8.5%, Fig. 3A, Fig. S3A). As before, we found that correct identification by ≥2 methods occurred in the majority of instances (Fig. 3A, red bars). The full list of target predictions generated by ≥2 methods for all 11 sRNAs (Fig. S3B) are included as Supplemental Dataset 1. On average the primary analysis by the pipeline took 1hr 15min ± 35min, using as many as 6 processing cores simultaneously. Re-collation of the results using different filters only took an average of 29s ± 6s.

**Figure. 3.**
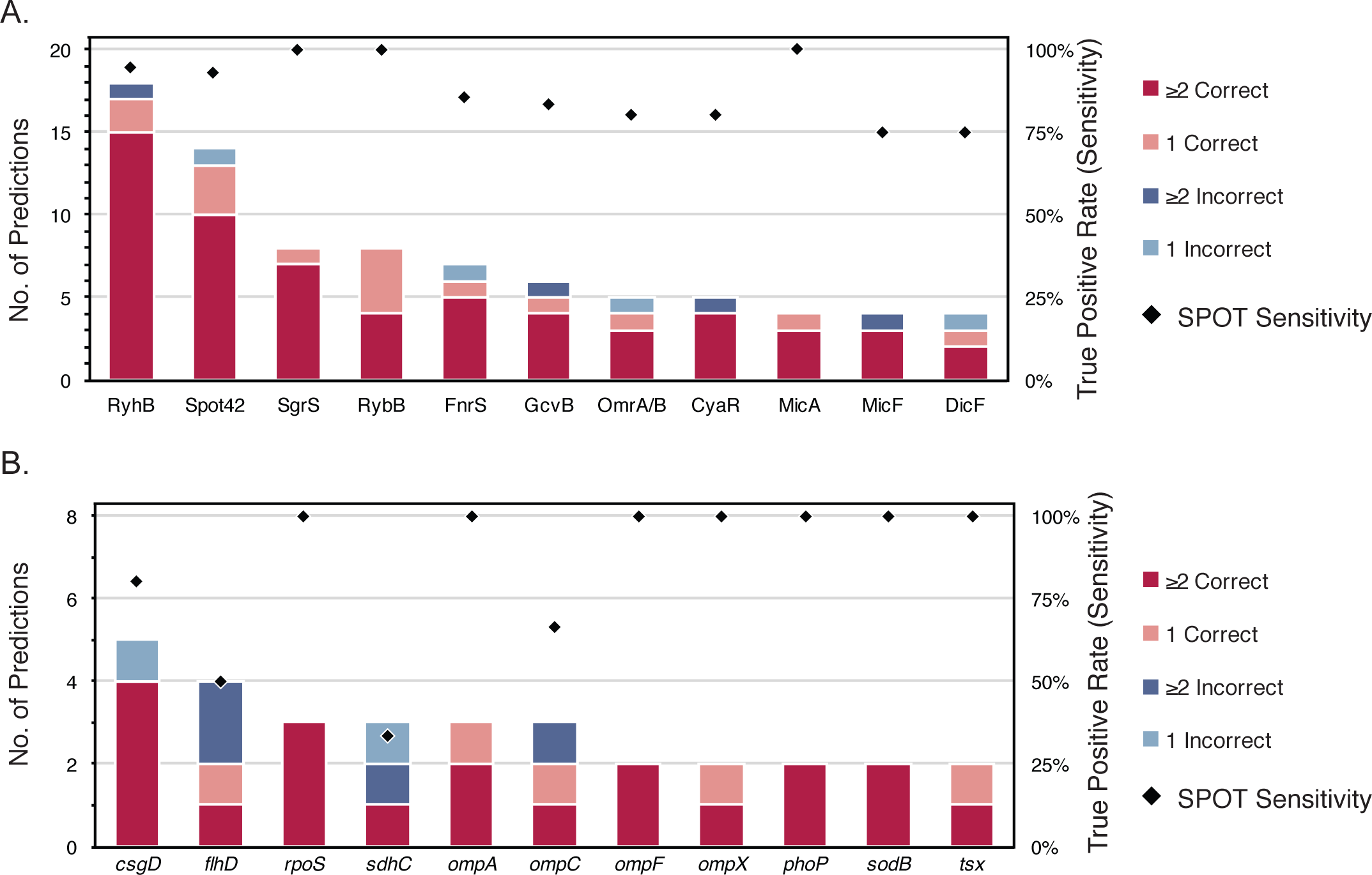
SPOT demonstrates high sensitivity for detecting targets of multiple sRNAs. (A) Along with RyhB and SgrS, nine additional sRNAs were analyzed with SPOT using the ‘Relaxed’ parameter set, demonstrating the robustness of the SPOT pipeline for correctly identifying sRNA-mRNA target interactions. Stacked bars show the number of experimentally validated mRNA targets correctly or incorrectly identified by 1 or ≥2 methods. Black diamonds indicate the overall True Positive Rate (sensitivity) of SPOT for each sRNA. (B) Eleven UTRs that are experimentally validated to interact with multiple sRNAs were used in a ‘reverse’ search in SPOT (i.e., using the UTR as the query and the sRNAs as the targets). The average sensitivity of this method is lower than in A., however, this is a novel means for identifying sRNAs that might affect genes of interest. Plots are drawn as in A.

### Extended application of the SPOT pipeline – mRNA as query sequence

The four individual algorithms are intended to identify the interaction of an sRNA with mRNA targets. However, a user may be interested in determining which known sRNAs interact with a specific mRNA of interest. Normally this would require running an individual search for each of the 10s to 100s of sRNAs from that organism. As part of our pipeline we have designed a feature that allows a user to input a custom annotation file for their reference genome. Therefore, instead of providing the list of mRNA targets, sRNAs can be provided to the algorithm and the relevant mRNA sequence, e.g., a 5′ untranslated region (UTR) of interest can be used as the query. We carried out this “reverse” analysis on 11 *E. coli*5′ UTRs that have already been demonstrated to interact with ≥2 different sRNAs. The results are comparable to the analysis using sRNAs as targets – known sRNA interactions were identified with an average sensitivity of 85% ± 24% (Fig. 3B). Moreover, using the 2-algorithm cutoff we were able to use this approach to predict 5 to 14 additional sRNAs that putatively bind the UTRs and could affect their regulation (Supplemental Dataset 1). We note that due to technical constraints the reverse search method can only be used with TargetRNA2, sTarPicker, and IntaRNA at this time. This approach is a novel feature that will facilitate ongoing sRNA research.

### Examination of novel RydC target predictions

We next sought to use the SPOT pipeline to identify additional targets for the poorly characterized sRNA RydC. RydC was reported to repress *yejA* mRNA (encoding an uncharacterized ABC transporter (43)) and *csgD* mRNA (encoding the master regulator of curli biogenesis (44)), but the molecular mechanisms of RydC-mediated repression were not reported. Fröhlich, *et al.*, (2013) demonstrated that RydC activates *cfa* mRNA, encoding cyclopropane fatty acid synthase. This activation involves RydC-dependent protection of *cfa* mRNA from RNase E-mediated degradation (3). Despite identification of these targets, the physiological function of RydC remains unclear. We used SPOT to identify additional targets of RydC as a means to gain further insight into its physiological role in *E. coli*.

Our strategy for RydC target identification was to combine computational and experimental data to generate an experimentally tractable list of putative targets for further validation. Experimental identification of putative targets was accomplished by pulse expression of RydC from an inducible promoter followed by identification of RydC-dependent changes in gene expression by RNA-seq. Vector control and P_*lac*_-*rydC* plasmids were maintained in a Δ*rydC* host strain grown in rich medium (LB) at 37°C. Expression of *rydC* was induced by addition of IPTG to cultures, and total RNA harvested at 10 minutes after induction. RNA-seq data output fastq files were analyzed with Rockhopper and exported as .xls files (Supplemental Dataset 2).

To identify putative RydC targets, the SPOT pipeline was applied to RydC using both stringent and relaxed parameters, with the former being more restrictive for window size and algorithm thresholds as described above. Similar to analyses for SgrS and RyhB, the relaxed parameters yielded a greater number of predictions than the stringent parameters. Potential targets that were predicted by ≥ 3 algorithms with the relaxed parameters are shown in Fig. 4 above the bold line. The RydC binding site for a validated target, *cfa* mRNA, was correctly predicted by 3 algorithms in the relaxed run. TargetRNA2 predicted a binding site that was inconsistent with the known binding site. The *cfa* prediction was absent in the stringent run, since the base pairing interaction between RydC and *cfa* mRNA takes place outside the window specified in the stringent run (Fig. 4). Some of the putative targets predicted by ≥ 3 algorithms were also differentially expressed in RydC pulse expression RNA-seq experiments (indicated under “Fold Change,” Fig. 4). Another set of genes were predicted as targets by ≥ 2 algorithms, and differentially expressed in RNA-seq experiments (Fig. 4, see targets below the bold line). Genes chosen for further analysis are listed in Table 1, along with information about their functions, differential expression in RNA-seq, predicted binding interactions, and algorithm predictions. Several other genes that did not meet the criteria for inclusion in Fig. 4 were also chosen for analysis because they had been described previously as RydC targets or because they encode proteins belonging to functional categories related to known RydC targets (Table 1).

**Table 1.**
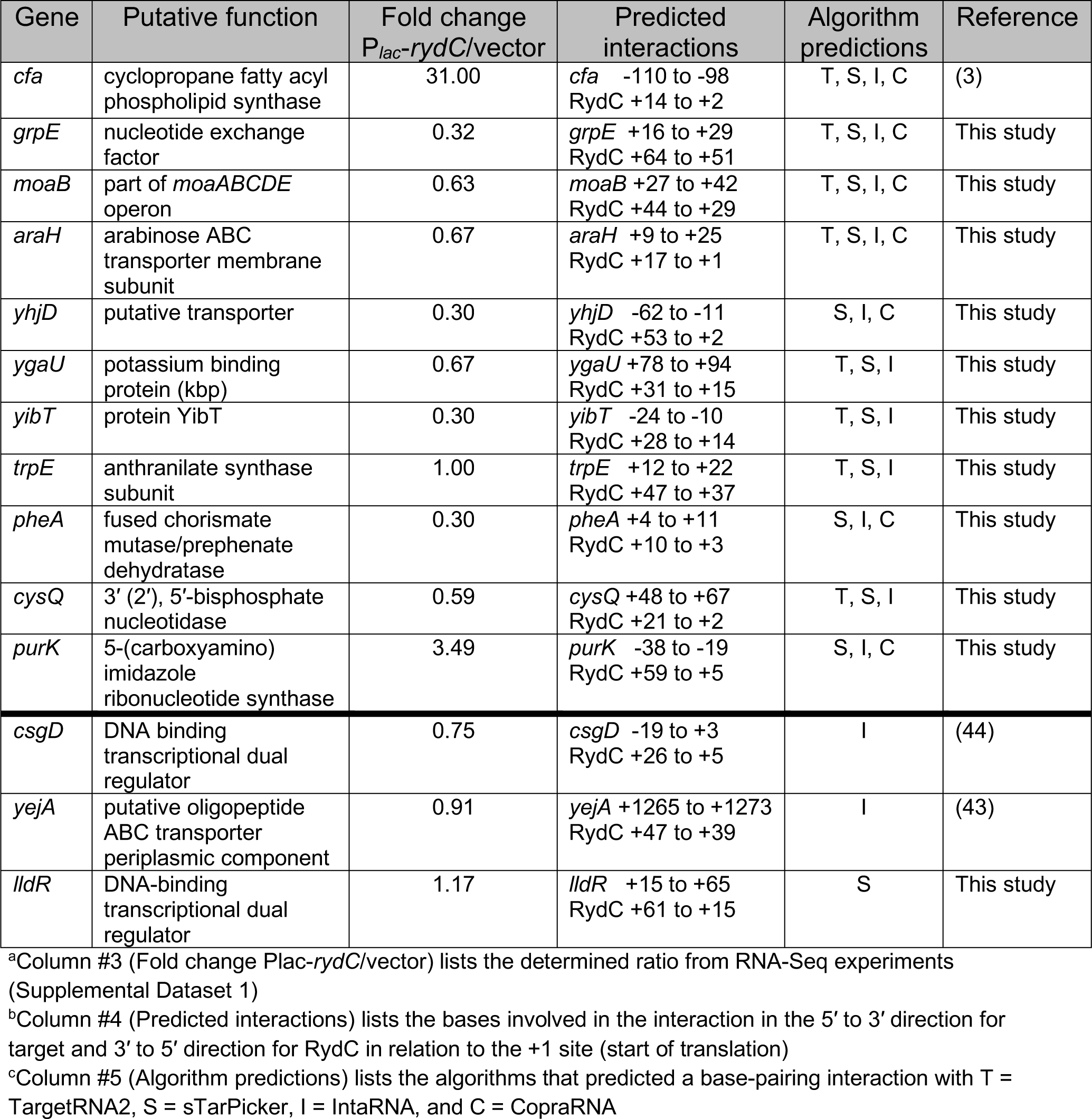
List of putative RydC targets chosen for further testing^a, b, c^.

**Figure. 4.**
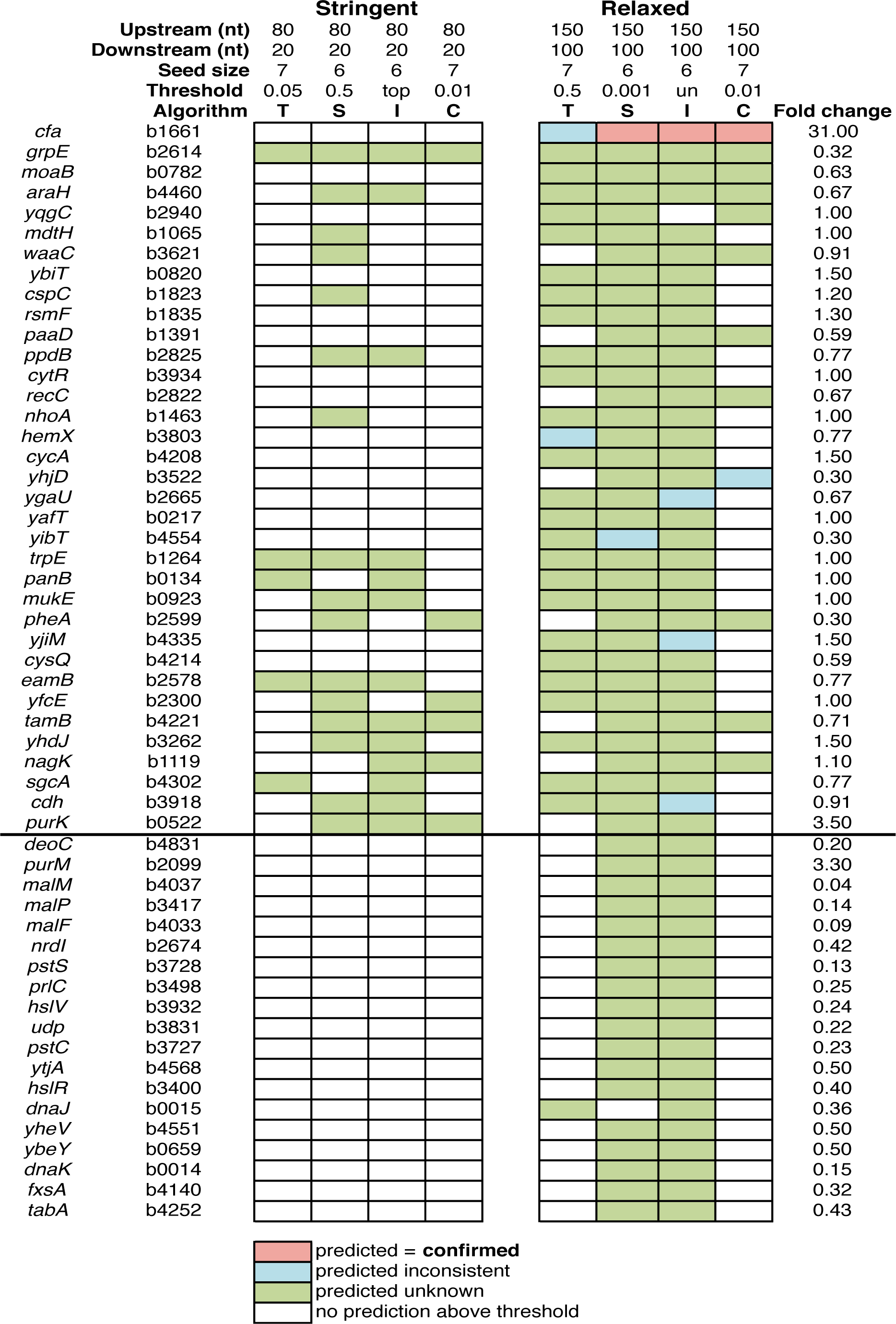
SPOT predictions for the sRNA RydC. Analyses were run with optimal seed sizes as determined in Fig. 2. Genes above the bold line denote those with ≥ 3 computational predictions, while genes below the line had 2 computational predictions and differential RNA-seq expression (fold change of ≥ 1.5 or ≤ 0.5, q-value of ≤ 0.005). Correctly predicted interactions for RydC are shown as pink cells, unknown predictions that were consistent among algorithms are shown in green, inconsistent predictions are shown in blue, and empty cells did not have any predictions above the indicated thresholds. Algorithms are abbreviated T=TargetRNA2, S=Starpicker, I=IntaRNA, and C=CopraRNA.

### Testing pipeline predictions for RydC

To test the targets selected for further validation for regulation by RydC, we constructed translational fusions to putative targets. These fusions were placed under the control of an arabinose-inducible promoter (P_BAD_) to eliminate possible indirect transcriptional effects. For each target, the entire 5′ UTR and part of the coding sequence (length variable, depending on the location of the predicted RydC binding site) was fused to *’lacZ* (Fig. 5A). Strains containing the reporter fusions were transformed with vector control or P_*lac*_-*rydC* plasmids and reporter activity measured after induction with IPTG.

**Figure. 5.**
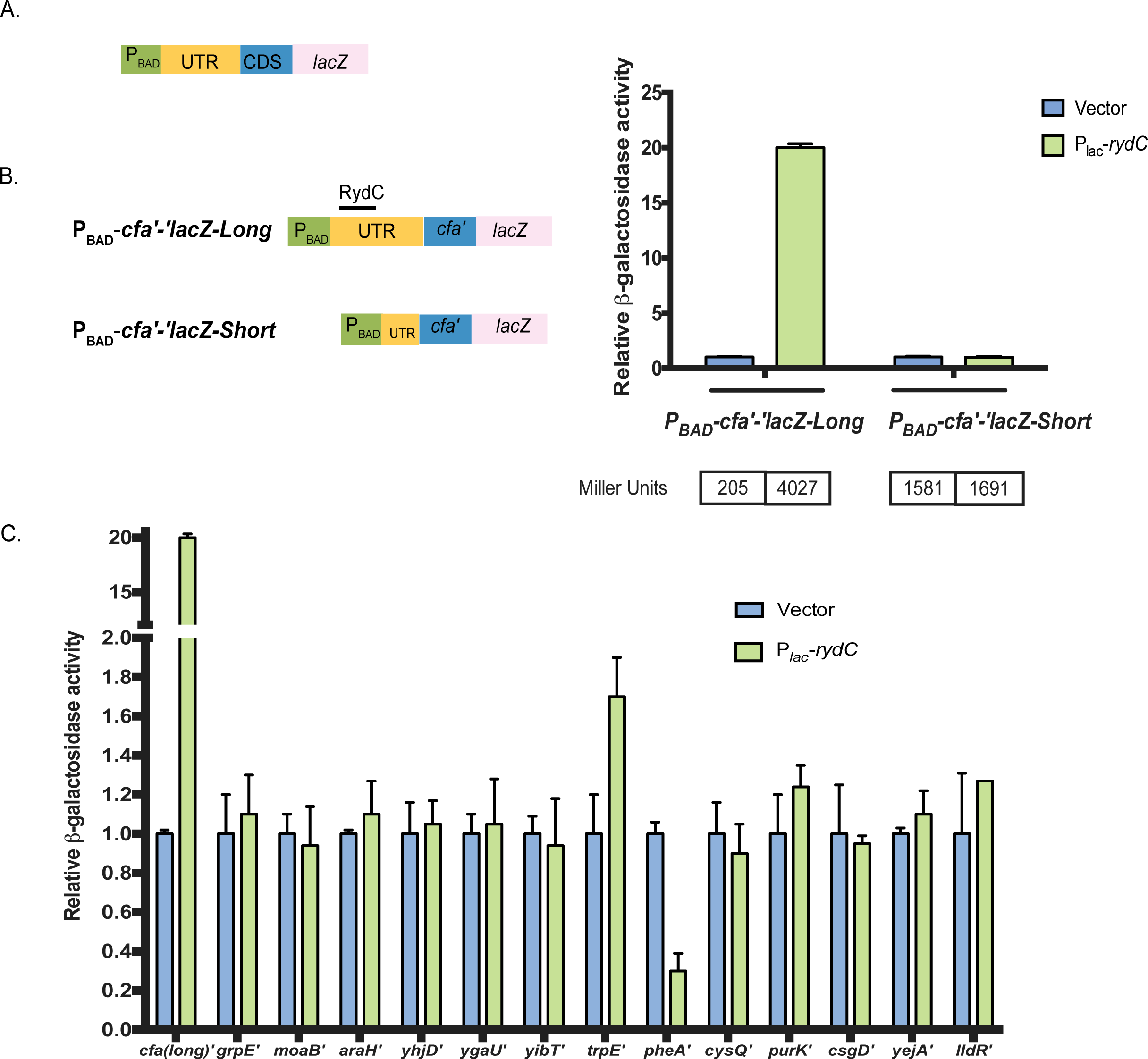
Validation of RydC target predictions. (A) The design for the translational*lacZ* constructs is shown, where the green box indicates the arabinose promoter (P_BAD_), yellow the untranslated region (UTR), blue the coding sequence (CDS), and pink the *lacZ* gene. (B) To confirm *cfa* as a RydC target, both full-length and shortened*cfa′-′lacZ* translational fusions were tested in backgrounds with vector or P_lac_-*rydC* plasmids. Expression of the reporter fusion was induced with 0.002% L-arabinose while induction of RydC was achieved with 0.1 mM IPTG. The activities were normalized to vector control and plotted as relative activity. The Specific Activity in Miller units are presented underneath the graph. These experiments were conducted as three independent trials with three biological replicates per trial. Error bars represent standard deviation among biological replicates from a representative trial. (C) Empty vector or RydC was overexpressed in strains with reporter fusion as indicated above. Expression of the fusion and RydC was induced as previously described. As a comparison, the positive control *cfa(long)’* was included in the experiment. These experiments were conducted as three independent trials with three biological replicates per trial. Error bars represent standard deviation among biological replicates from a representative trial.

In *Salmonella,* RydC was demonstrated to activate *cfa* translation by occluding an RNase E cleavage site to stabilize the *cfa* mRNA (3). Conservation of RydC-*cfa* mRNA interactions between *E. coli* and *Salmonella* as well as SPOT identification of *cfa* as a putative RydC target (Fig. 4, Table 1) suggest that *E. coli* RydC regulates *cfa* in a similar manner. To confirm this, we constructed two translational fusions: P_BAD_-*cfa’-’lacZ*-Long, which contains the RydC binding site, and P_BAD_-*cfa’-’lacZ*-Short, which lacks the RydC binding site (Fig. 5B). RydC production strongly activated the long fusion, increasing activity by >20-fold compared to the vector control strain (Fig. 5B). As expected, activity of the short fusion lacking the RydC binding site was unaffected upon RydC induction (Fig. 5B). These results support the model that *cfa* mRNA is a directly regulated by RydC in both *S. enterica* and *E. coli*.

Strains harboring reporter fusions to 13 other putative targets (listed in Table 1) were transformed with vector control and P_*lac*_-*rydC* plasmids and β-galactosidase assays were performed after a period of RydC induction (Fig. 5C). Only two of the target fusions were differentially regulated by the criteria we selected (≥1.5-fold or £0.5-fold) in RydC-expressing cells compared to the vector control (Fig. 5C). These two targets were *pheA* and *trpE*, which both encode proteins involved in aromatic amino acid biosynthesis. Previous studies (43, 44) reported RydC-dependent translational repression of the *yejA* and *csgD* mRNAs, though we note that specific and direct base pairing interactions with RydC were not demonstrated. Our translational fusions to these putative targets did not show any differential regulation in response to RydC expression (Fig. 5C).

### RydC regulates genes in aromatic amino acid biosynthetic pathways

In RNA-Seq experiments, levels of *pheA* mRNA were reduced to ~30% of control levels when RydC was ectopically expressed (Supplemental Dataset 2). Likewise, in RydC-producing cells, activity of the P_BAD_-*pheA’-’lacZ* fusion was ~30% that of the vector control (Fig. 5C). The predicted RydC-*pheA* mRNA base pairing interaction involves the 5′ end of RydC and the coding region of *pheA,* directly adjacent to the start codon (Fig. 6A). The P_BAD_-*pheA*’-’*lacZ* fusion encompasses all of the 5′ UTR and 645-nt of the coding region. A reporter derived from this has mutations that disrupt the predicted base pairing with RydC, resulting in the P_BAD_-*pheA67*’-’*lacZ* fusion with mutations G9C/G10C (Fig. 6A). A *rydC* allele with compensatory mutations (C4G/C5G) was constructed and named RydC5. The mutations in RydC5 abrogated regulation of the wild-type P_BAD_-*pheA*’-’*lacZ* fusion. Likewise, the mutations in P_BAD_-*pheA67*’-’*lacZ* prevented regulation by wild-type RydC. The compensatory mutant pair: P_BAD_-*pheA67*’-’*lacZ* and RydC5 had restored regulation, albeit not to fully wild-type levels. Together, these data suggest that RydC targets *pheA* mRNA for translational repression. Due to the location of the base pairing interaction in the translation initiation region, mechanism is likely direct occlusion of ribosome binding to *pheA* mRNA by RydC.

**Figure. 6.**
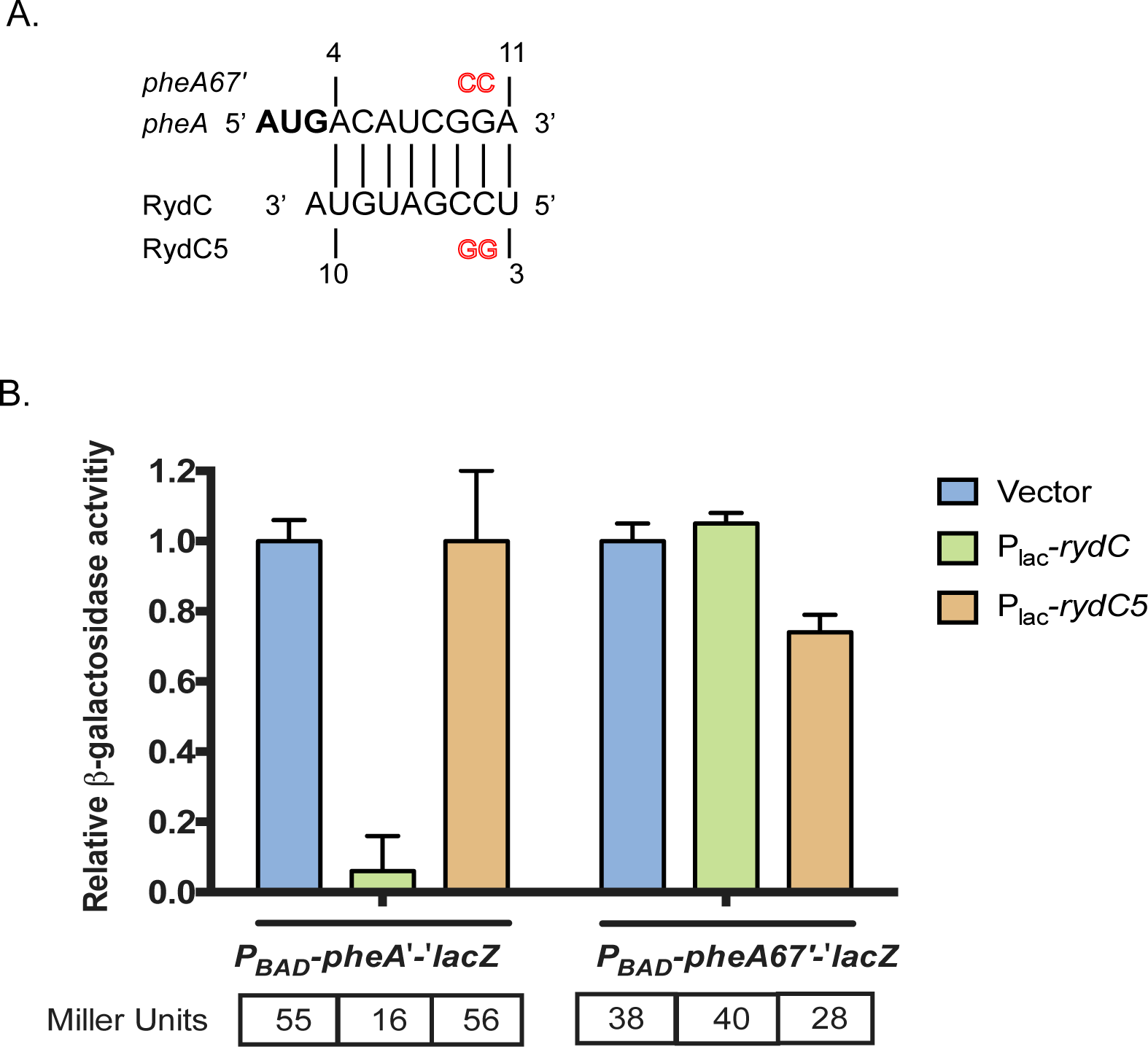
RydC represses *pheA* translation. (A) The predicted base pairing between *pheA* mRNA and RydC from IntaRNA. The residues highlighted in red represent point mutations made for each of the variant fusions/RydC alleles. The numbers are in relation to the +1 of RydC and the AUG of *pheA*. To test *pheA* as a putative target, both full-length and mutated (*pheA67′*) *pheA′-′lacZ* translational fusions were tested. (B) RydC or a RydC variant (RydC5) were overexpressed in the *pheA′* and *pheA67′ lacZ* fusion backgrounds. Expression of the fusion and RydC was induced as described for Fig. 5. The activities were normalized to vector control and plotted as relative activity. The data were analyzed and reported as described for Fig. 5.

Another new putative RydC target is *trpE,* which encodes a component of the anthranilate synthase involved in tryptophan biosynthesis. A P_BAD_-*trpE’*-*’lacZ* fusion encompassing the 30-nt *trpE* mRNA 5′ UTR and 42-nt of *trpE* coding sequence was activated upon RydC production by slightly less than 2-fold (Fig. 5C, 7B). The predicted RydC-*trpE* mRNA base pairing interaction involves sequences near the 3’ end of RydC and sequences within the *trpE* coding sequence. Point mutations in the *trpE* reporter fusion (C20G/C22G) resulted in the mutant reporter P_BAD_-*trpE20’-’lacZ,* which was not substantially upregulated when wild-type RydC was produced (Fig. 7B). Because of the unusual pseudoknot structure of RydC (3, 44) mutations in the 3’ end of RydC have a dramatic impact on RydC stability (45), thus we were not able to test a RydC compensatory mutant that would restore pairing to the *trpE20* mutant fusion. However, we did construct a second *trpE* fusion, P_BAD_-*trunc-trpE’-’lacZ,* which was truncated to remove the putative RydC binding site (Fig. 7B). This fusion was no longer activated by RydC at all. These observations suggest that sequences early in the *trpE* coding sequence are important for RydC-mediated increase in *trpE* translation.

**Figure. 7.**
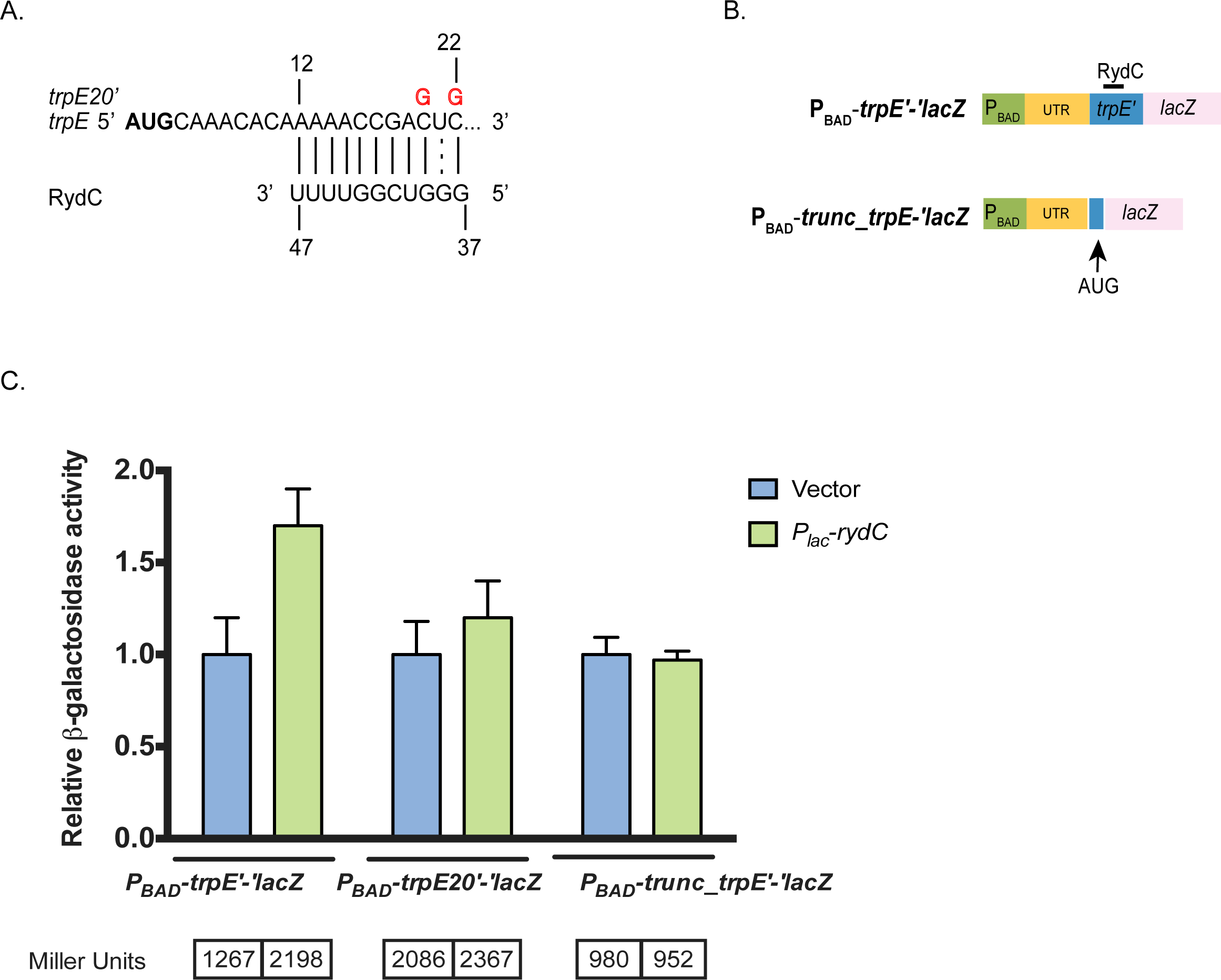
RydC activates *trpE* translation. (A) The predicted base pairing between *trpE* mRNA and RydC from IntaRNA. The vertical/dotted lines represent the seed region for base pairing interactions. The residues highlighted in red represent point mutations made for each of the variant fusions/RydC alleles. The numbers are in relation to the +1 site of RydC and the ‘AUG’ of *trpE*. To test *trpE* as a putative target, both full-length, mutated (*trpE20′*), and truncated (*trunc_trpE′*) *trpE′-′lacZ* translational fusions were tested. (B) Empty vector or RydC were overexpressed in the *trpE′*, *trpE20′*, and *trunc_trpE′ lacZ* fusion backgrounds. Expression of the fusion and/or RydC was induced as previously described. The experiments were conducted and data analyzed and presented as described for Fig. 5.

## DISCUSSION

Over the years, many sRNAs have been discovered and characterized using both computational and experimental methods. Although target discovery of sRNAs still remains the rate-limiting step in sRNA characterization, many new techniques have been developed to overcome that obstacle. Some techniques take a purely computational approach to target prediction, including the target-prediction algorithms we have included in SPOT, (9, 10, 11, 12, 13) and others we have not included (47-57). Experimental techniques to identify bacterial sRNA targets have also expanded. Many of these use affinity purification or co-immunoprecipitation approaches, with or without crosslinking (15, 17, 20, 58, 59, 60). To help streamline the process of sRNA target identification, the SPOT pipeline was constructed to be used in conjunction with other identification methods. In this study, we showed that the SPOT pipeline achieved ≥ 75% sensitivity and ≤ 50% false positive rate when at least 2 methods converged on a prediction for the well-characterized sRNAs SgrS and RyhB (Fig. 2D-E). Expanding our analysis to other bacterial sRNAs, we found that the pipeline sensitivity was equal to or exceeded that of any individual method (average = 84% ± 8.5%, Fig. 3A, Fig. S2). As before, we found that correct identification by ≥2 methods occurred in the majority of instances (Fig. 3A). Furthermore, SPOT can be applied to the reverse situation where a user can search for potential sRNAs that regulate their UTR of interest. We found through these analyses that for 11 *E. coli*5’ UTRs with ≥2 known interactions with sRNAs, the analysis gave an average sensitivity of 85% ± 24% (Fig. 3A).

To test the utility of SPOT in identifying novel sRNA-mRNA target interactions, we used it to predict targets of the poorly characterized sRNA RydC, which had been described to regulate three genes: *yejA*(43), *cfa*(3), and *csgD* (44). Through SPOT analyses and filtering based on experimental data, we generated a list of putative RydC targets (Table 1). Reassuringly, SPOT identified the true RydC target, *cfa* mRNA, and correctly predicted the known binding site on this target (Table 1, Supplemental Dataset 1). The other two reported targets, *yejA* and *csgD,* were not identified by the SPOT computational pipeline, nor were these genes differentially regulated in our RydC pulse-expression RNA-seq analyses (Supplemental Dataset 2). Since no specific direct binding interactions were shown for RydC-*yejA* or RydC-*csgD*, we postulate that the previously observed regulation of these targets by RydC may be indirect. The SPOT pipeline also correctly identified 2 additional RydC targets, *pheA* and *trpE* (Table 1, Figs. 5C, 6, 7). RydC represses *pheA* translation, likely by a mechanism common to repressing sRNAs. Binding of RydC to sequences around the Shine-Dalgarno region would prevent ribosome binding and inhibit translation initiation. The mechanism of RydC-dependent activation of *trpE* appears to be more complex. The *trpE* gene is part of the *trpLEDCBA* operon responsible for L-tryptophan biosynthesis, which is regulated by both the *trpR* repressor and an attenuation mechanism (46). Depending on the availability of L-tryptophan, the ribosome can either stalls at or moves quickly through Trp codons in the *trpL* ORF. When Trp is abundant, the ribosome rapidly completes translation of *trpL,* which prevents co-transcriptional formation of an antiterminator hairpin and allows formation of a transcription terminator just upstream of the *trpE* coding sequence. When Trp is limiting, ribosome stalling at the Trp codons allows formation of an antiterminator structure, which promotes transcription elongation into downstream Trp biosynthesis structural genes. While sequences within the *trpE* coding sequence have not been implicated in the Trp-dependent attenuation mechanism, it is possible that the sequences including the RydC binding site are responsible for yet another layer of regulation of these genes, perhaps at the level of translation. Alternatively, sequences in the *trpE* coding sequence could have long-range interactions with the upstream terminator or antiterminator sequences and RydC binding could modulate those interactions.

Our study and evaluation of a combinatorial approach to identify mRNA targets of sRNAs of interest represents a step toward accelerating a rate-limiting step in sRNA characterization. The SPOT pipeline is able to streamline the process of running individual algorithms, which can take hours to days, by reducing the run times significantly for all 4 algorithms at once (under 2 hours). Since the pipeline runs all 4 algorithms simultaneously, a more narrowed down, comprehensive list is generated, negating the need for manually selecting targets from individual algorithm runs. However, every method has drawbacks and though SPOT is a powerful tool, it has limitations as well. For instance, a 50% false positive rate (the average for well-characterized sRNAs analyzed in this study) is still high even though it is markedly better than the false positive rates of predictions made by any single algorithm. As experimental approaches for sRNA-mRNA target identification continue to improve, the power and accuracy of SPOT’s combinatorial approach to sRNA-target binding site predictions will likewise improve. Another factor impacting the accurate prediction of sRNA binding sites by SPOT is the user-defined search window. The majority of early examples of sRNA-mediated regulation involved sRNAs binding in translation initiation regions of target mRNAs. Thus, most existing sRNA target prediction algorithms have default windows set to search around start codons. As more sRNA-mRNA interactions are validated and mechanisms of regulation studied, we and others have found increasing numbers of examples of sRNA-mRNA interactions that occur outside this window. Some of these interactions are primary or only interactions responsible for sRNA-mediated regulation of the mRNA, e.g., RydC-*cfa* mRNA (3), SgrS-*yigL* mRNA (39), which both involve mRNA sequences far upstream of the start codons. Yet other interactions involving mRNA sequences far from translation initiation regions represent secondary or auxiliary binding interactions that nevertheless play important roles in regulation (18, 38).

For the sRNA SgrS, there are two binding sites for its interaction with *asd* mRNA (18), but SPOT was only able to predict the primary binding site. We expect that there are other examples where the algorithms have failed to identify alternate or additional binding sites. This is currently an area of development and once implemented, will serve as a valuable asset in identifying putative targets for a sRNA of interest.

Taken together, the combinatorial approach revealed two new targets, *pheA* and *trpE*, in the RydC regulon. Interestingly, both PheA and TrpE are involved in the chorismate metabolic pathway, with PheA using chorismate as a substrate in L-Tyrosine/L-Phenylalanine biosynthesis and TrpE for L-Tryptophan biosynthesis. Interestingly, RydC repressed *pheA* whereas it activated *trpE,* an unusual case since both are involved in amino acid biosynthesis in divergent pathways. In the case for *trpE,* the mechanism of positive regulation is unique in that the base pairing interaction takes place 12-22 nt downstream of the start codon. RydC could possibly serve as a sRNA modulator of the biosynthetic pools of amino acids by activating/repressing *trpE*/ *pheA* mRNA expression when necessary. As an aside, chorismate is also a substrate for production of the *E. coli* siderophore enterobactin, which is synthesized under iron limiting conditions. Mutations in *fur*, *tyrA*, *pheA*, or *pheU* resulted in increased enterobactin production since the chorismate pools were used for enterobactin synthesis (62). These observations suggest that there may be conditions where RydC impacts the iron starvation stress response, perhaps forming a regulatory network that intersects with that of the well-characterized iron starvation stress response sRNA, RyhB. To better understand these potential connections, future work will be aimed at characterizing the regulators and conditions controlling synthesis of RydC.

With the implementation of the SPOT pipeline, combined with RNA-Seq and MAPS data, we were able to add to the RydC regulon and expand its network. Whether this regulatory network is exhaustive remains to be determined. We note that there were other RydC-mRNA binding interactions predicted by SPOT that were not analyzed further here. Moreover, there are additional sRNA-mRNA interactions predicted by SPOT for the other sRNAs that were run through the pipeline (Supplementary Dataset 1) and it is likely that more bona fide interactions are among those predictions. All in all, we developed a streamlined method for sRNA-mRNA binding site predictions that leverages the strengths of many pre-existing algorithms. We showed the robustness of SPOT for identification of true sRNA-mRNA interactions using well-characterized and poorly characterized sRNAs. We anticipate that SPOT will become a valuable tool for many investigators who have found interesting sRNAs and wish to identify potential mRNA targets for further characterization.

## Acknowledgements

We thank members of the Vanderpool lab, past and present, the laboratory of Dr. James Slauch and members of AMK’s committee, Drs. Cronan, Metcalf and Slauch for constructive feedback throughout the course of the project. We are grateful to Dr. Eric Massé for sharing information about RyhB targets, and to Dr. Hanah Margalit for information about putative RydC-target interactions based on experimental RIL-seq data. We are very appreciative of Dr. Alvaro Hernandez and staff at the Roy J. Carver Biotechnology Center for assistance with RNA-seq.

## Funding

Funding for this work was provided by the National Institutes of Health (R01 GM092830) to CKV and supported AMK. The Roy J. Carver Charitable trust award (#15-4501) to PHD, and University of Illinois at Urbana-Champaign and University of California Riverside start-up funds to PHD also supported the work.

